# Targeted Sequencing of Pancreatic Ductal Adenocarcinoma Tissue

**DOI:** 10.1101/2023.01.13.523431

**Authors:** Kiruthikah Thailli, Christian Benzing, Alberto Quaglia, Debashis Sarker, Claire M Wells

**Affiliations:** School of Cancer and Pharmaceutical Sciences, Kings College London., London UK

## Abstract

Pancreatic ductal adenocarcinoma (PDAC) is an aggressive cancer and the majority of patients present with metastatic disease. Metastatic spread requires a reorganisation of the actin cytoskeleton, a process that is dependent on the Rho family GTPases and their interaction with subsequent downstream effectors. The p21 activated kinases (also known as the PAKs) are effectors of Rho GTPases Cdc42 and Rac and play key roles in cell migration and survival. PAK4 is overexpressed in PDAC and can reciprocally activate the PI3K pathway. Hepatocyte growth factor (HGF), via c-Met, is an established activator of PAK4 and the PI3K pathway and may promote PDAC invasion. Next generation sequencing (NGS) was performed on 31 PDAC tissue using a pre-determined targeted panel to attempt to identify novel mutations in the PAK: PI3K pathway. Targeted NGS identified several recurrent mutations including 5 PAK4 mutations in PDAC tissue.

## Introduction

It is known that pancreatic ductal adenocarcinoma (PDAC) has a poor prognosis, with devastatingly low 5-year survival rates. Yet some patients remain cancer free more than 10 years after their surgery. There are clearly key differences in their biology that prevents recurrent or progressive disease that are yet to be determined. The answer undoubtedly lies within specific genetic patterns of their cancer and more extensive genomic interrogation may uncover such unique prognostic signatures. PDAC is known to have relatively low tumour mutational burdens compared with other malignancies on average 30 somatic mutations per tumour (Lawlor et al., 2021). This has often been cited as one of the reasons for the failure of immune-therapeutics when used alone. Yet of the mutations present, several have been identified as recurrently altered. Numerous studies have shown that *KRAS* mutations are present in nearly 100% of all samples, with common oncogenic events in *TP53, SMAD4* and *CDKN2A* also prevalent and several other genes have been reported at lower rates. Of particular interest is the p-21 activated serine/threonine kinase family (PAK1-6) which are reported to play a role in the progression of a number of different tumour types (King et al., 2014). PAK4 specifically has been associated with PDAC (King et al.,2017), but the PAK mutational landscape in PDAC has not been explored.

Recent years have seen rapid developments in next generation sequencing (NGS) technology enabling the cancer genome to be studied in its entirety. Deep sequencing of a whole genome can now be performed within a day compared to historical sanger-sequencing that required over a decade. At present, the required infrastructure to study whole genomes (such as computer capacity and bioinformatics support) means whole genome sequencing is still predominantly limited to the research setting rather than guiding medical management. The advent of targeted sequencing of individual or small clusters of genes has revolutionised personalised cancer care. In targeted sequencing, a subset of genes or regions of the genome are isolated determining small base changes (substitutions), insertions or deletions. In addition, targeted sequencing can also identify structural changes including copy number variations. Lung, skin and colon cancer management are now guided by the presence or absence of activated mutations which are routinely screened for at diagnosis.

## Materials and Methods

### Tissue collection and laser microdissection

Ethics approval was sought and granted to use tissue stored at King’s College Hospital (KCH) NHS Foundation Trust. Patients had consented prior to surgery for their pancreatic tissue to be stored within the KCH liver biobank. Ethics approval was granted to use this tissue for this specific project (IRAS 194304). Samples were fixed formalin paraffin embedded (FFPE) and were of pancreas that contained pancreatic ductal adenocarcinomas that had been resected at the time of surgery as treatment. Tissue not needed for clinical management was stored in the biobank. All samples were from local early-stage cancers. Specimens were identified and prepared by specialist hepatobiliary pathologists. 48 Samples were sliced and mounted on to RNAse free glass slides ready for tissue dissection, one layer was stained for haematoxylin and eosin (H and E) and cytokeratin 7 (CK7) to ease identification of cancer cells prior to laser micro-dissection. Cancer cells were identified by a specialist hepatobiliary pathologist. Samples were dissected using a laser microscope and tissue was collected into Eppendorfs. These were stored at room temperature until DNA extraction.

### DNA extraction and Sequencing

DNA was extracted using the Qiagen DNA kit. 48 Samples were placed in a microcentrifuge tube and 1ml xylene was added. The sample was vortexed and then centrifuged for 2 minutes. The supernatant was discharged. Ethanol was added, and the mix was centrifuged and the supernatant extracted. The tube was then kept open at room temperature for 10 minutes until the residual ethanol had evaporated. The remaining pellet was suspended in 180uL of buffer ATL (pre-prepared) and proteinase K. If the sample did not appear to be digested, this step was repeated.

The sample was then incubated at 56°C for 1 hour or until the sample had completely lysed. The sample was then incubated at 90°C for 1 hour, then centrifuged. Buffer AL was then added to the sample and mixed by vortex. A further 200uL of ethanol was added and the sample vortexed. The sample was then transferred to a QIAmp MinElute column and centrifuged at 6000g for 1 minute.

Buffer AW1 was added and the sample centrifuged again. Buffer AW1 was added and the sample centrifuged. The sample was then placed in a new tube and then the sample was eluted using a Mini Elute column following the addition of buffer ATE. Samples were checked for DNA quality using the nanodrop and 31 samples were taken forward for genomic analysis using the Illumina Miseq platform.

## Results

In this project, archival tissue taken at the time of surgery for curative surgery for early stage PDAC between 2000 and 2010 was used. A panel was designed to perform targeted sequencing of 25 separate genes using the Illumina MiSeq NGS platform **(Table 1)**. Genes were chosen following an extensive literature search.

**Table 1.**
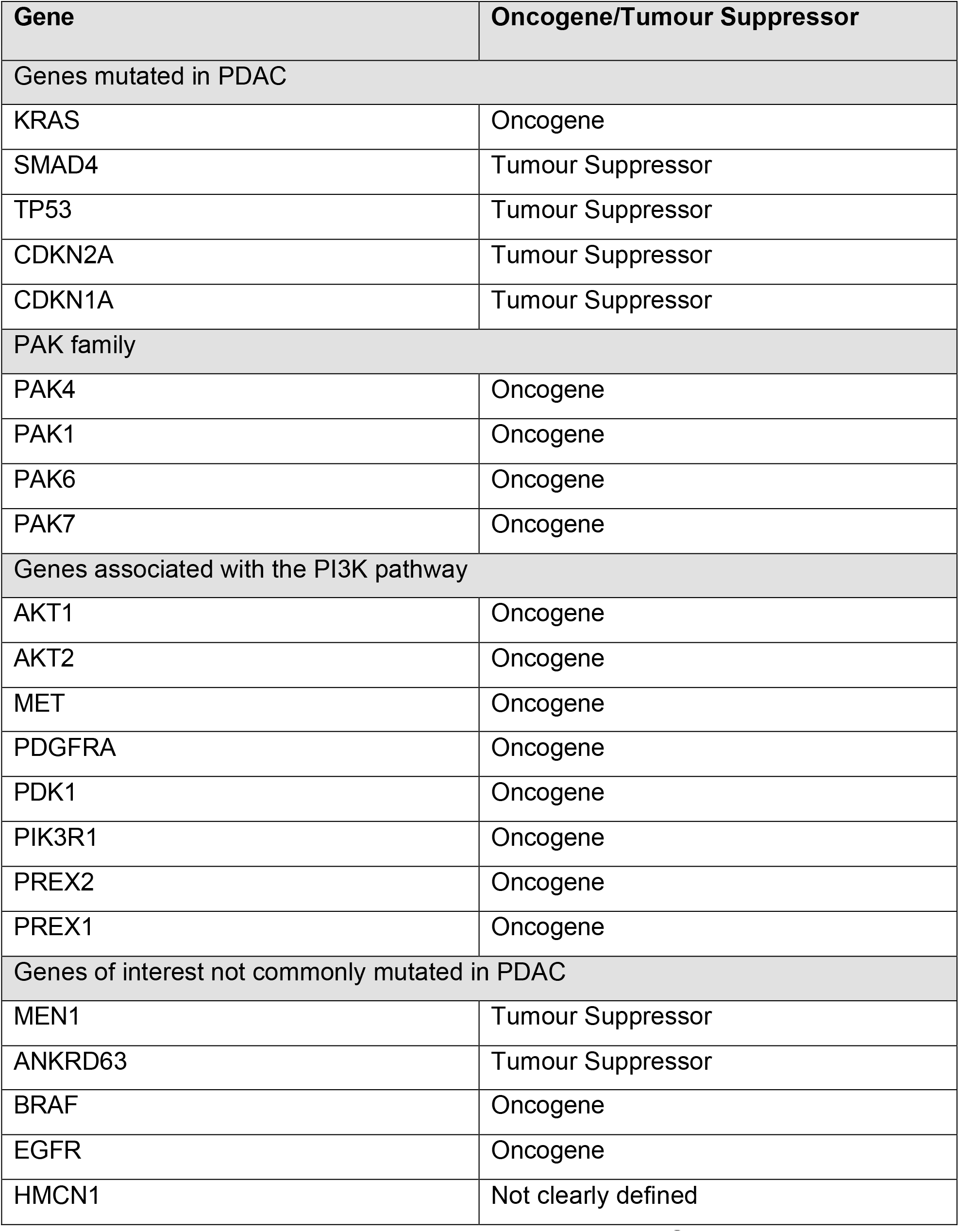
List of genes used in the targeted panel for Next Generation Sequencing. A combination of known oncogenes and tumour suppressor genes were used. Some genes, the mechanism in which they produce oncogenic affects remains undetermined

*KRAS, TP53, SMAD4* and *CDKN2A* are known to be altered in *PDAC. PAK1,4,6* and *5* were chosen as genes of interest due to their previous links with PDAC and our own studies associating their activity with cancer progression. Note *PAK5* is known as *PAK7* on the sequencing plate and thus *PAK5* will be referred to as PAK7 thereafter. *PAK4* mutations are rare but have been found to be significant (Whale et al., 2013) when identified and therefore were included for analysis as deep sequencing in PDAC samples, has not previously been performed. The remaining genes were chosen due to their role in the PI3K pathway (Thillai et al., 2017). These included *AKT1* and *AKT2, PIK3R1, PREX1* and 2. Other genes such as *EGFR* and *BRAF* are frequently mutated in other malignancies, but not commonly in PDAC but were included in the panel for assessment. Finally, genes that are not commonly mutated but are found in hereditary conditions such as *MEN* were included.

### Prevalence of mutations per patient

All 31 patients harboured mutations in 1 or more of the selected genes and the median number was 13 **(Figure 1)**. The smallest number of mutations was seen in patient 31, (3 mutations) and the highest was 20 (patient 12). To visualise the different mutations seen in the 31 samples by genes, a heat map was created. (**Figure 2)**. This includes the different variations of mutated genes. The most common mutation was in *PREX1* which was found in all samples, followed by the tumour suppressor gene *MEN1* which was found in 30 samples. Mutations were classified into categories (**Table 2**) and only group 1 and 2 categories (Group 1= HIGH IMPACT and Group 2=MODERATE impact) were further analysed.

**Table 2.** Categories used to classify genetic mutations. Only categories 1 and 2 are likely to have a clinical impact.

|  |
| --- |
| **Category I HIGH IMPACT** |
| Known in the literature to be clinically significant and causative<br>Evidence for the pathogenicity in the locus specific databases as being causative Is associated with disease in the GWAS catalogue<br>Is associated with a tumour site in COSMIC Is validated in clinical study in NCBI SNP |
| **Category II MODERATE IMPACT** |
| Introduction of a stop codon In-frame exon deletion<br>Mutates the initiation codon (ATG)<br>Missense mutation of the normal stop codon<br>Annotated probably pathogenic in NCBI SNP database<br>Deletes nucleotide(s) that lead(s) to a shift of reading frame Deletes exon which results in shift of reading frame |
| **Category III LOW IMPACT** |
| Modifies UTR 3'<br>Is non-synonymous coding variant in start Modifies UTR 5'<br>Results in codon change Results in codon insertion<br>Results in codon change and codon insertion Results in codon change and codon deletion Deletes UTR 5' |
| **Category IV** |
| Is synonymous coding variant in stop<br>Is synonymous coding variant in start<br>Is synonymous coding variant<br>Is intergenic<br>Is intronic variant<br>Unlikely to produce a cryptic splice site |
| **Category V** |
| Allele origin is somatic<br>Allele origin is germline<br>Allele origin is unknown<br>Is labelled suspect in NCBI SNP |

**Figure 1.**
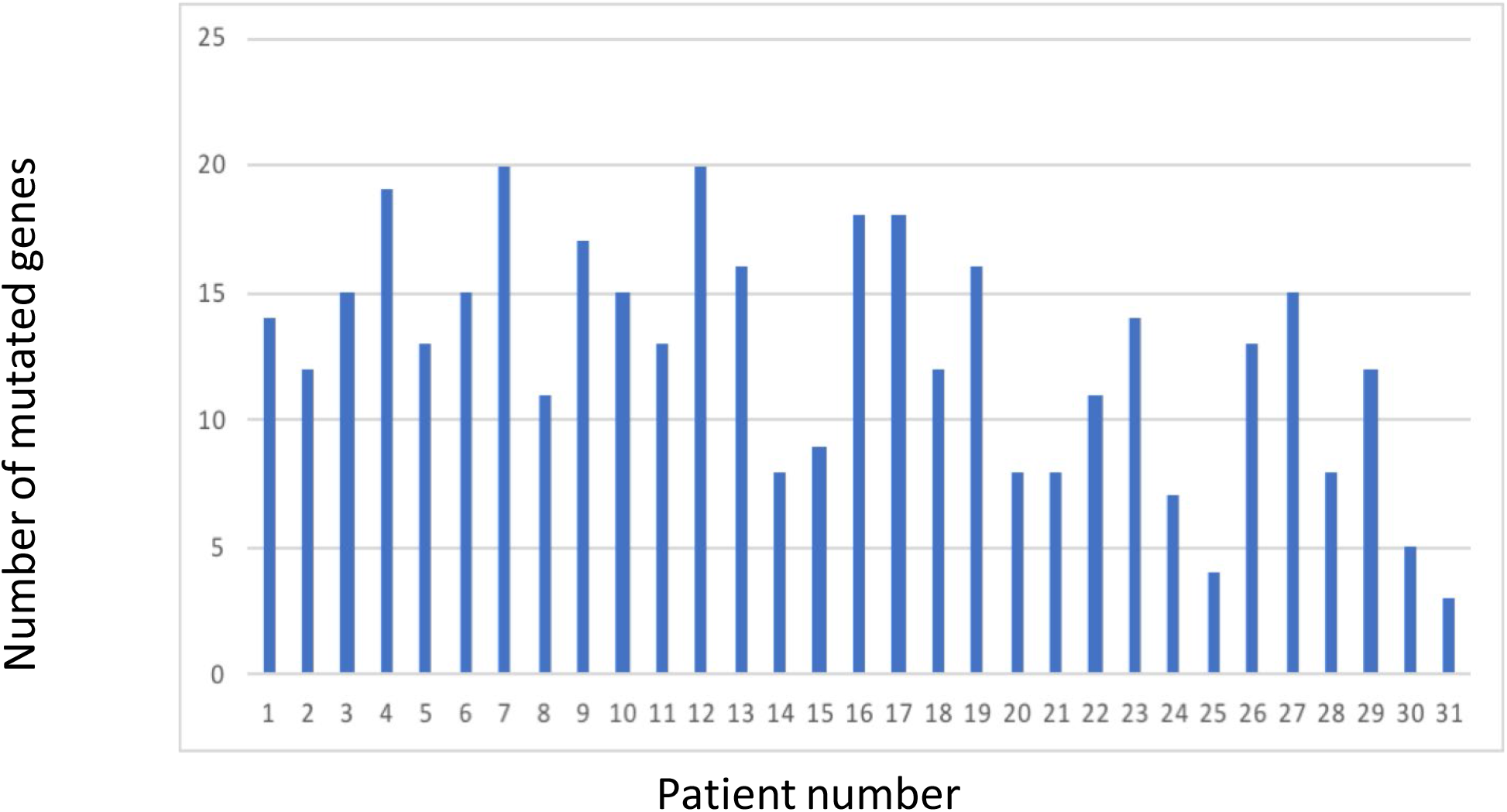
Graph of mutations across 31 different patient samples. Targeted sequencing revealed all 31 samples harboured mutations. The highest number was 20, present in 2 samples.

**Figure 2.**
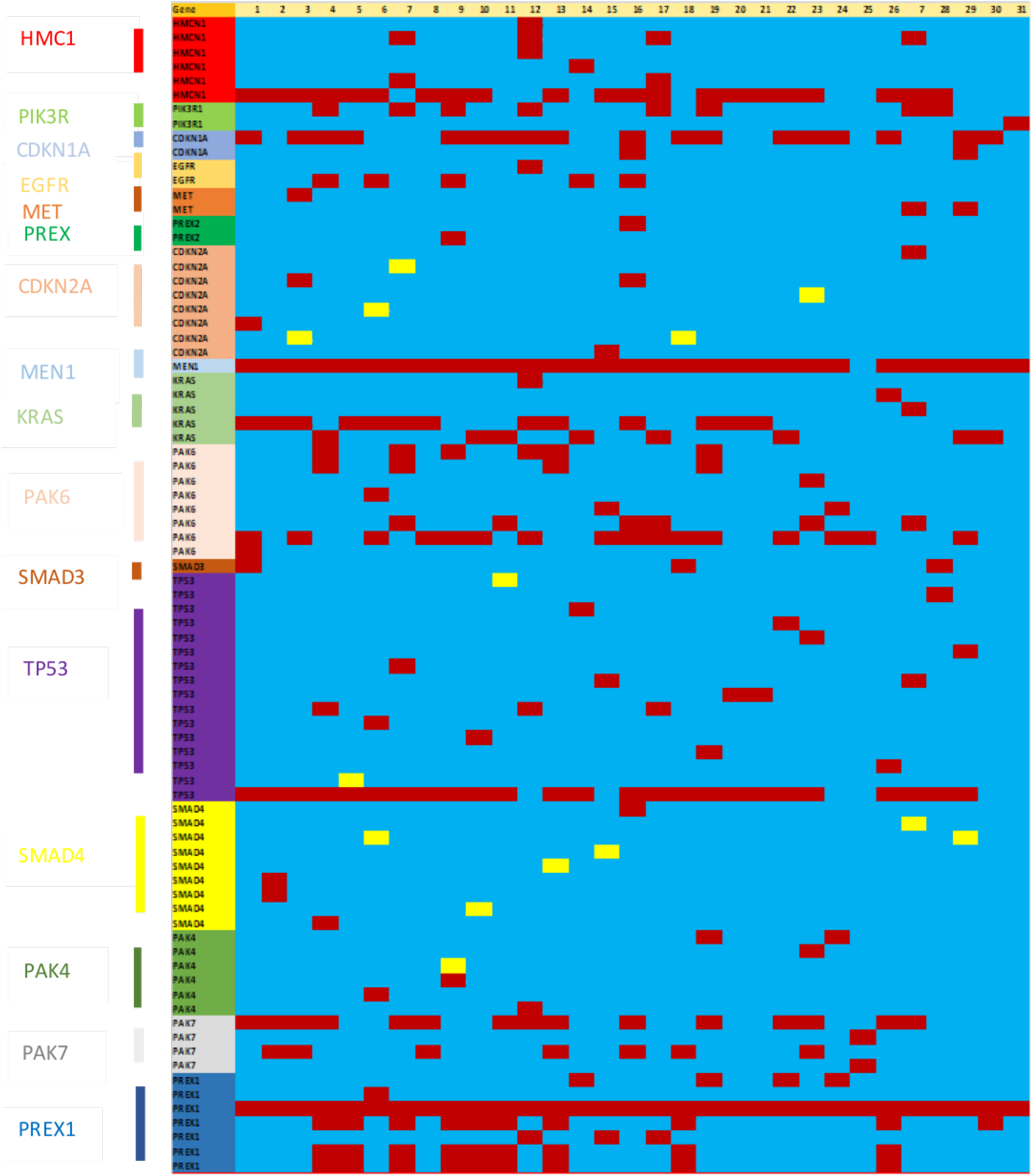
Heat map of mutated genes across 31 different patient samples. Yellow indicates HIGH impact mutations. Red indicates MODERATE impact mutations.

### Overview of mutations

In the 31 patients, 19 mutations were category 1. These included 7 frame shift mutations, 3 splice site mutations, 1 stop lost mutation and 3 stop gain mutations. Genes affected were *PAK4, SMAD4, TP53, CDKN2A, PDK1* and *HMNC1* **(Table 3)**. There were 81 group 2 non-synonymous coding mutations present in the 31 patient samples **(Table 4)**. These included 18 different genes, *HMCN1, PDK1, PDGFRA, PIK3R1, CDKN1A, EGFR, MET, PREX2, CDKN2A, MEN1, KRAS, PAK6, SMAD4, TP53, SMAD3, PAK4, PAK7 and PREX1*. The most prevalent was a T/C substitution in *PREX1* at chromosome 1:47253150 which was found in all 31 samples. 6 patients had a *PAK4* mutation (2 were A/T substitutions at the loci 19: 39663768).

**Table 3.**
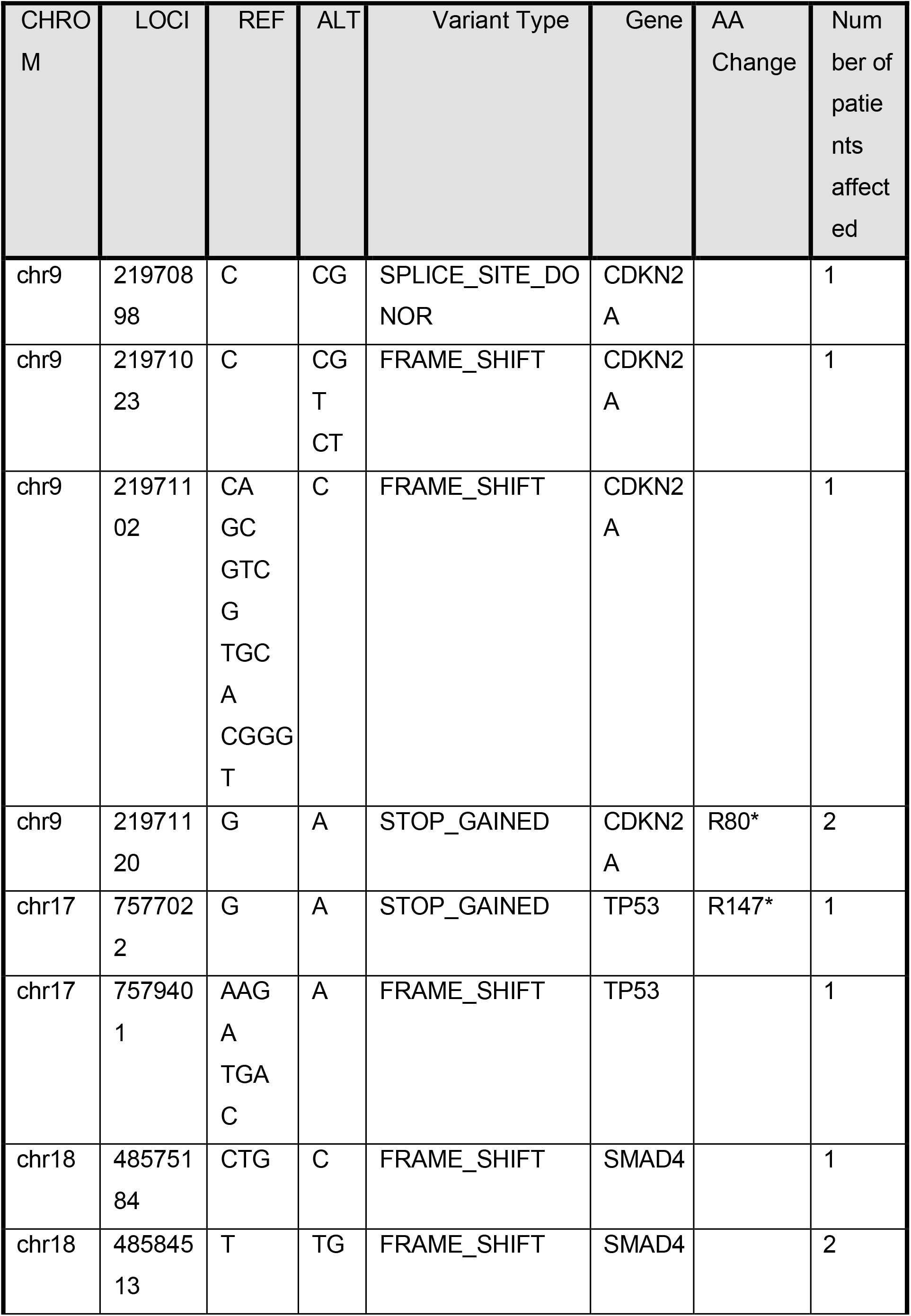
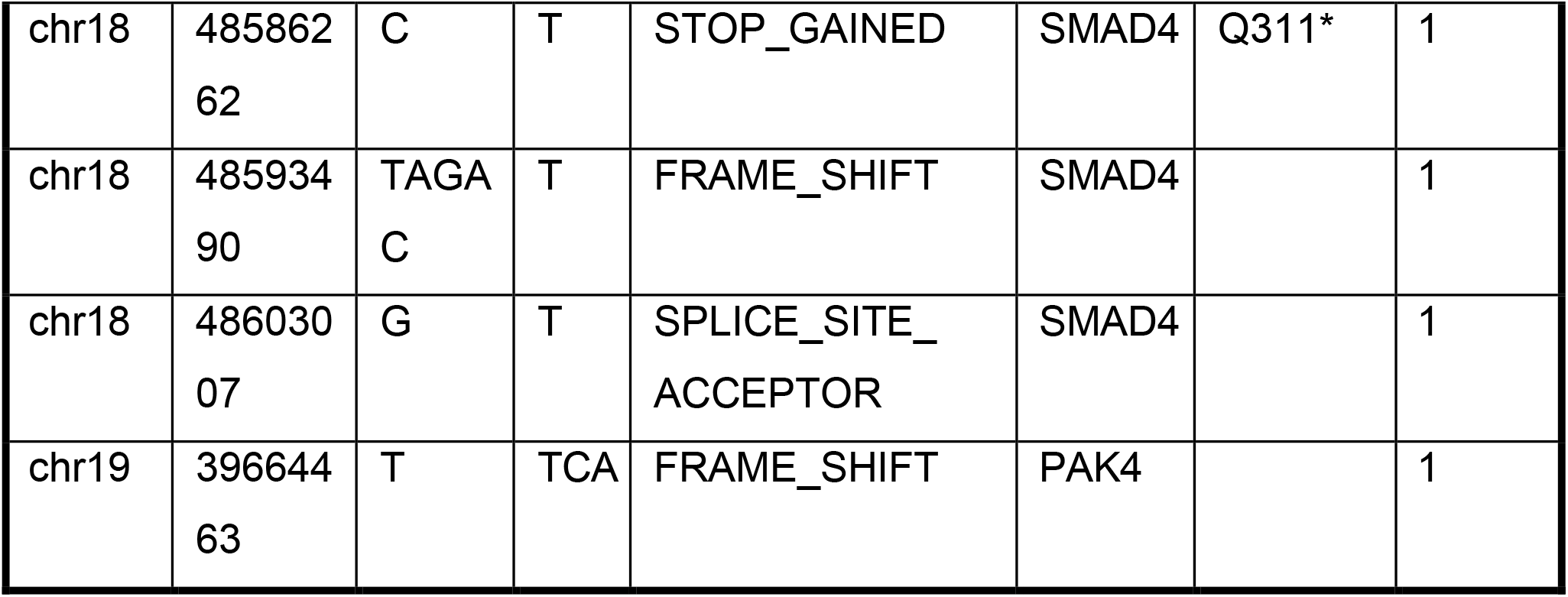
A list of all Category 1 (HIGH impact) mutations. These included, frame shift, stop codon and splice site mutations. Chr =chromosome, REF=reference, ALT=alteration

**Table 4.**
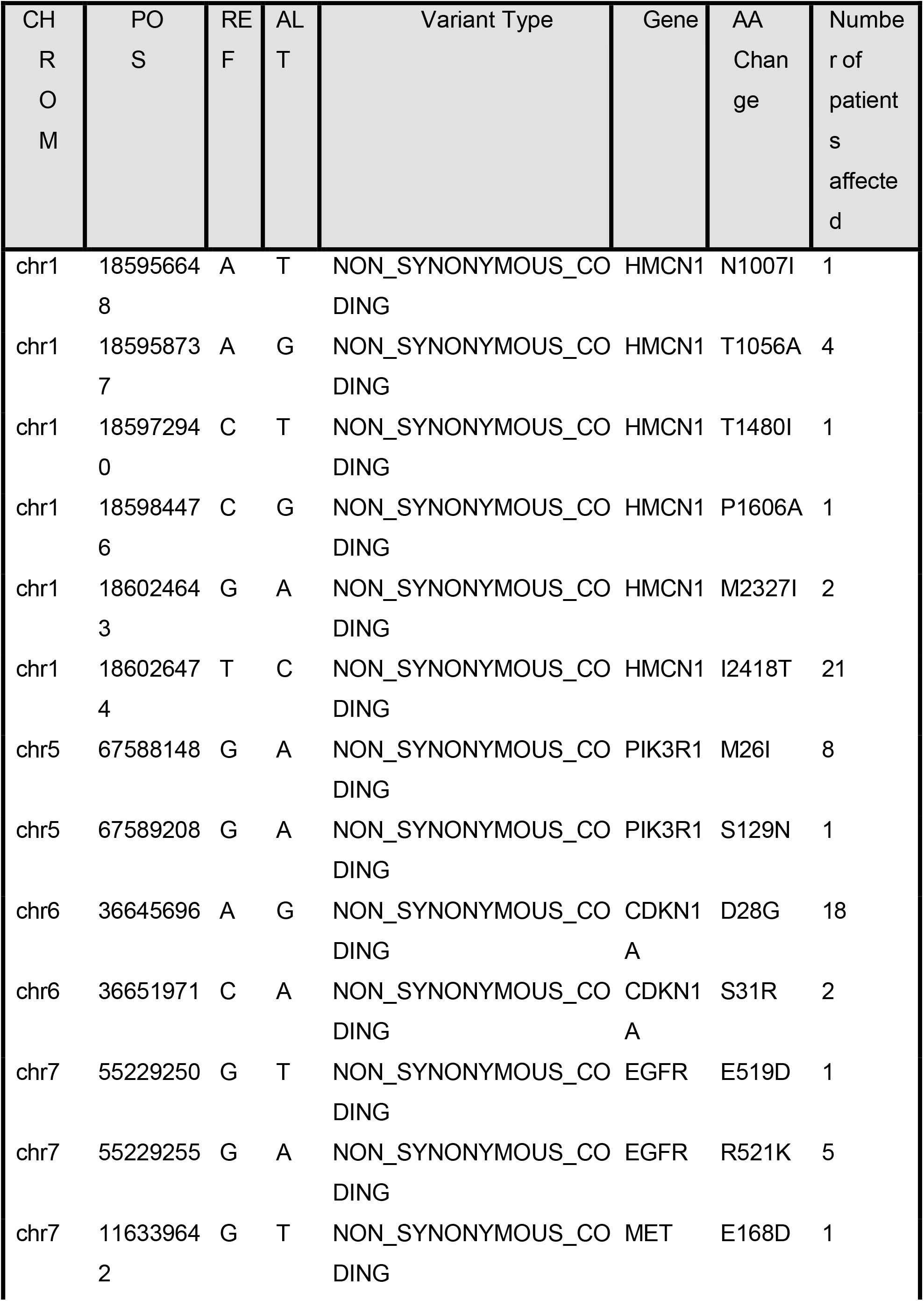
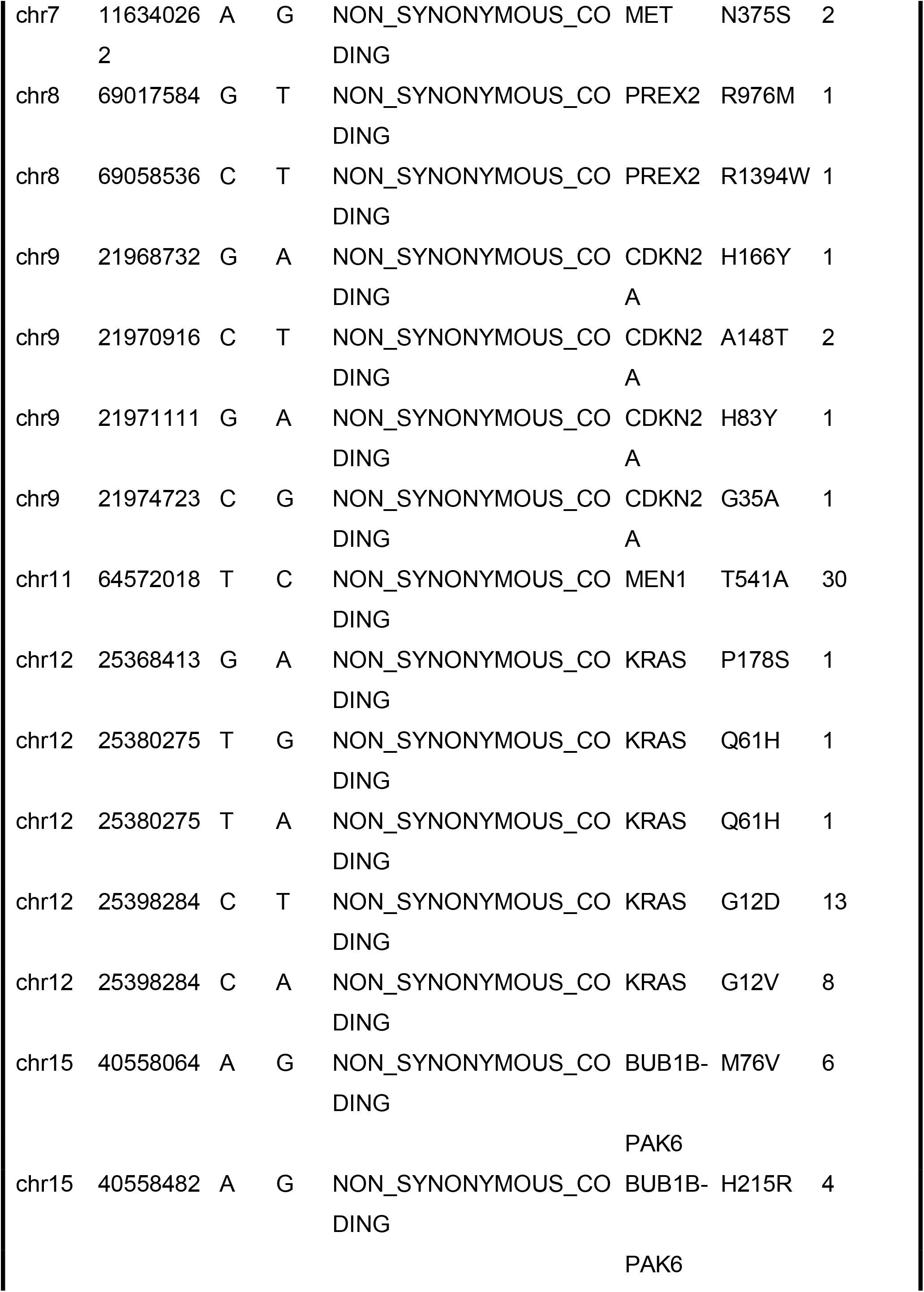
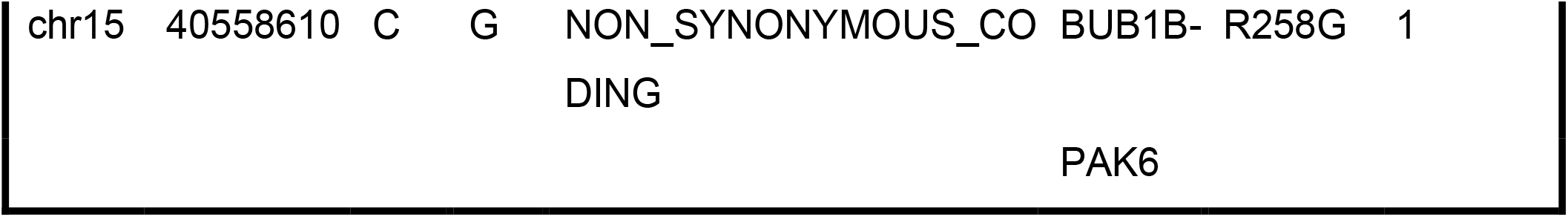
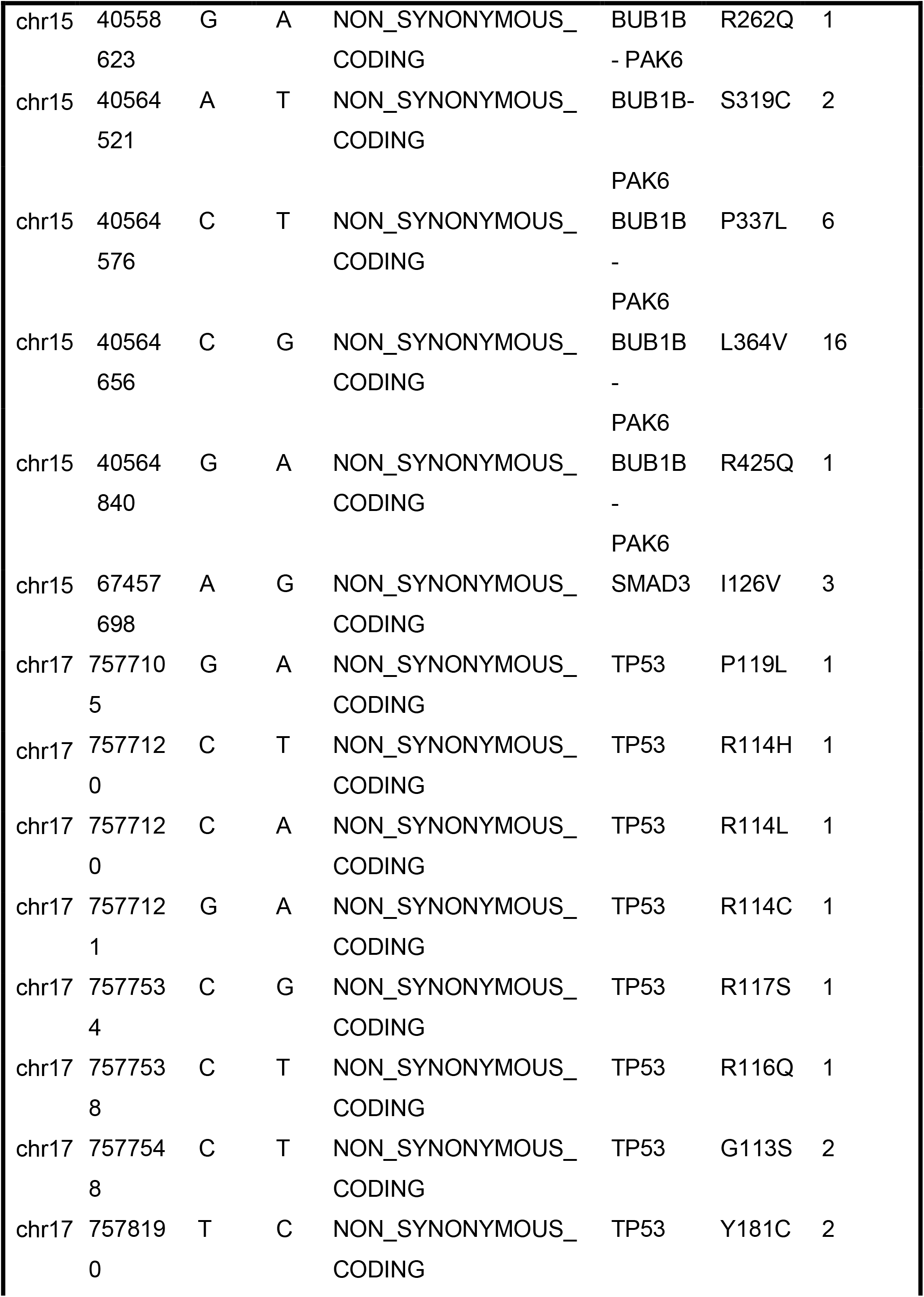
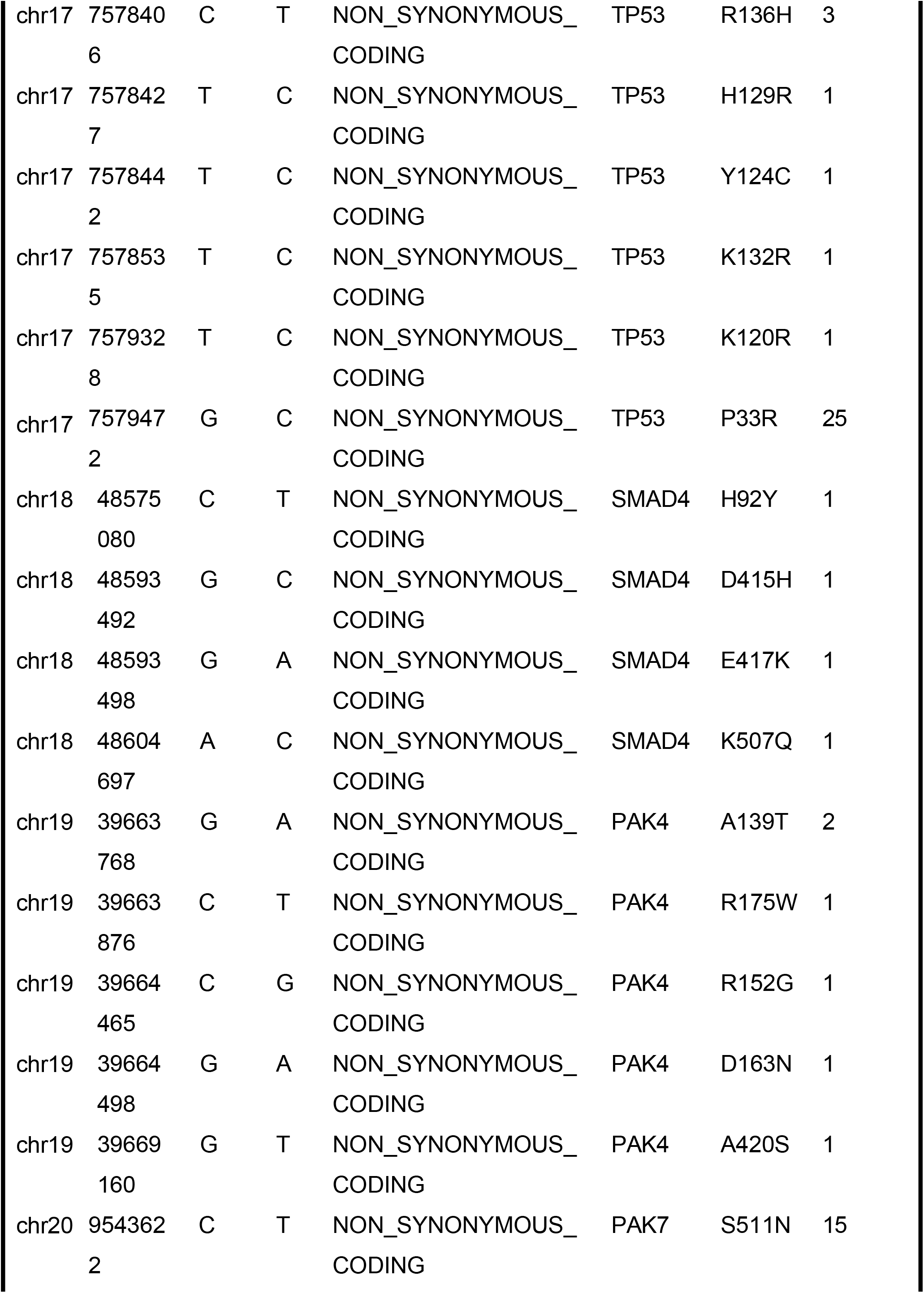
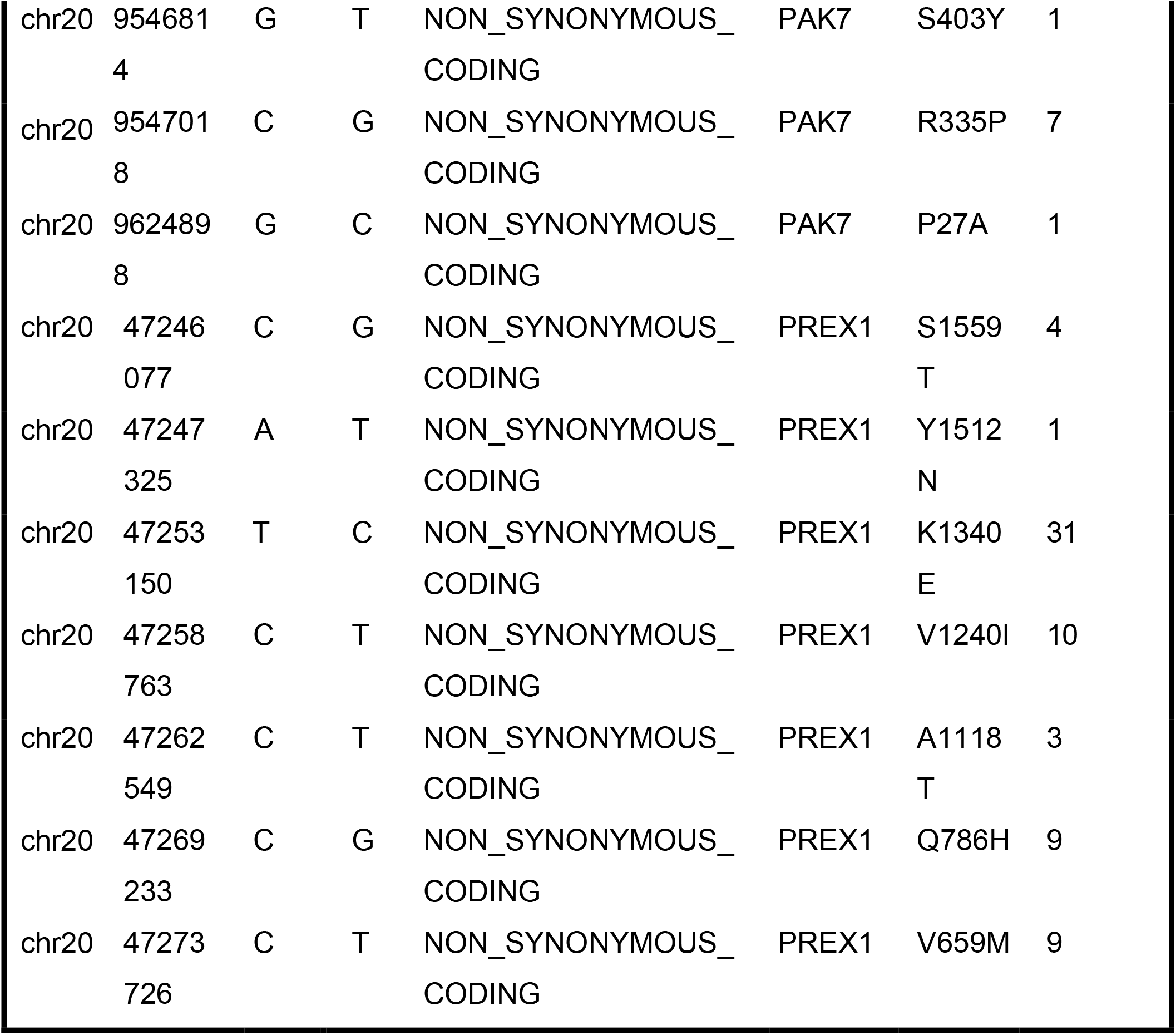
A list of all Category 2 (MODERATE impact) mutations

### KRAS mutations

Nearly all PDAC tumours are expected to harbour a *KRAS* mutation, this is often associated with pre-neoplastic events and is the first mutation to occur. 24/31 (77%) of patients harboured a mutation in *KRAS*. The most prevalent was the G12D at 12:25398284, which is a change from a Glycine (hydrophilic non-charged) to an Aspartate (negatively charged) acidic amino acid. 8 patients had a mutation in G12V at 12: 25398284, which is the change from a glycine at position 12 to a valine (hydrophobic, non-polar) amino acid. 2 patients had a Q16H mutation at 12: 25380275, whereby a glutamine (acidic negatively charged) at position 61 changed to a histidine (basic, positively charged). Finally, a patient had a p178S change at 12: 25368413 from a proline (non-polar hydrophobic) to a serine (polar non hydrophilic). This mutation was a modification of the UTR5’ resulting in the insertion of a codon insertion.

### TP53 mutations

25 patients had a *P33R* mutation, which is a common missense mutation that changes proline (non polar hydrophobic) to arginine (basic and positively charged). Other individual mutations were found, including a stop-gain mutation at 17: 7579472, position with R147, which has previously been reported in the literature.

### CDKN2A mutations

5 patients had a CDKN2A mutation. These included 5 non-synonymous coding mutations, 2 at 9: 21970916.

### SMAD mutations

SMAD4 was found in 9 patients, with 5 category 1 mutations. 3 frame shift mutations, 4 nonsynonymous coding, 1 stop gained and 1 splice site acceptor. For example, the category 1 frame shift mutation seen in the 4th row of table, 18:48584513T>TG affects transcription of SMAD4 are affected. The frame shift variant has a HIGH impact. For SMAD3, 3 mutations were found in 3 different patients. These were non-synonymous coding mutations at 1126V and all of MODERATE impact.

### PIK3R1 mutations

PIK3R1, mutations have strongly been implicated in cancer, especially breast and have been proposed as predictive markers for therapeutics to the PI3K pathway. 8 patients had a nonsynonymous coding, with a change from methionine to isoleucine. 1 patient had a serine to an arginine. All had a MODERATE impact.

### EGFR mutations

EGFR mutations are uncommon in PDAC. 6 patients had an EGFR mutation. 5 patients had a non-synonymous coding mutation in position 519 where glutamate was changed to an aspartate. 1 patient had a point mutation in position 521 where an arginine was changed to a lysine. Both were of MODERATE impact.

### CDKN1A mutations

18 patients had a non-synonymous coding variation at position 28 where aspartate was changed to a glycine and 2 patients had a change from serine to asparginine. Both of these mutations had a MODERATE impact.

### HMNC1 mutations

*HNMC1* mutation was found in 20 patients, with several patients harbouring more than one aberration. The most common non-synonymous coding mutation was found in position 2893 and 2418. Known variation at 1007, 1480, 2837, 2808 were found as well as HIGH impact splice site acceptor.

### MET mutations

The *MET* gene is the proto-oncogene that codes for *CMET*. MODERATE impact mutations were found at position 7: 116339642 G/T and 7: 116340262 A/G.

### MEN mutations

*MEN* is not commonly mutated in PDAC samples and is associated with *MEN* type 1 which is commonly linked with endocrine tumours. 30 patients had a mutation that was MODERATE at 11: 64572018 T/C.

### PREX mutations

3 patients had a serine changed to a proline at position 478 resulting in a non-synonymous coding change in *PREX1*, 31 patients had a K1340E (lysine changed to glutamate). Further changes seen at position 124, 659 and 786. *PREX2* displayed 2 counts of non-synonymous coding. The Variant is MODERATE impact. This results in codon change of Cgg/Tgg resulting in an amino acid change of R1394W and a further change at R976M.

### PAK mutations

15 patients had a non-synonymous *PAK7* coding mutation at 20:9546814. The Variant is MODERATE impact. This results in codon change of a Gc/aAc and Amino acid change of S511N. Further MODERATE mutations were seen in 20:9543622, 20:9657018 and 20:9624898. One HIGH impact *PAK4* mutation was identified **(Table 5)**. This was a frame shift identified in *PAK4* at the loci 19:39664463, T>TCA. 5 patients had a non-synonymous coding mutation of PAK4. To understand the prevalence and importance of *PAK4* mutations compared with the reference genome, non-synonymous coding variations were inputted into available databases. Ensemble, 1000 genome, and ExAC was used. 12 instances of a variation at the loci 19:39663768 G>A (A139T) were found. This included liver cancer, ovarian cancer, lymphoma and benign conditions. Using an online software tool SIFT –that sorts the tolerant and intolerant amino acid substitutions to predict the phenotypic effect.

**Table 5.**
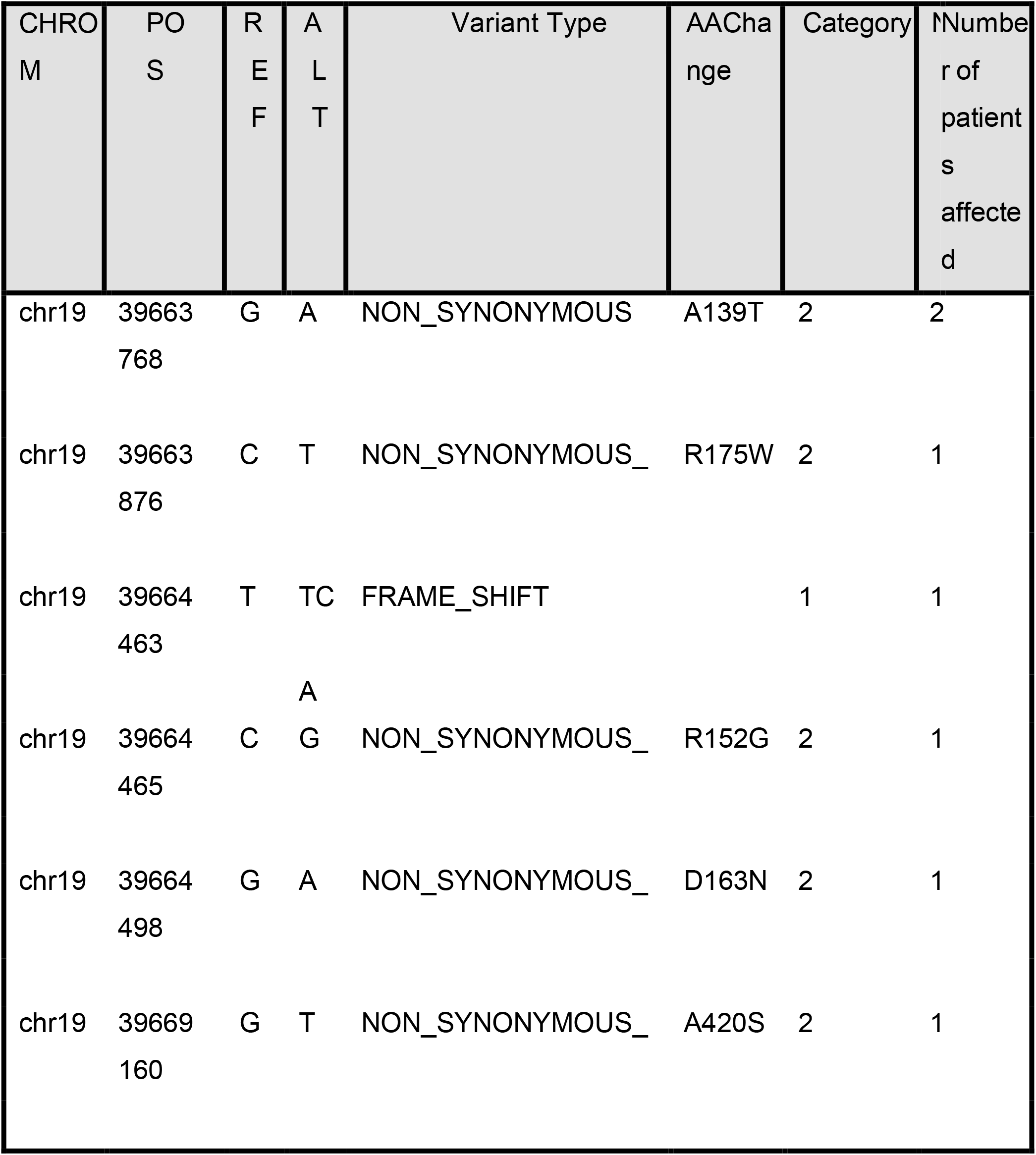
PAK4 mutations

A139T delivered a score of 0.53 suggesting it would be tolerated, rather than deleterious. A change at 19:39664465 was also predicted to be tolerated as was 19:39664498 and 19:39669160. Again with reference to the A139T mutation, a separate measure (STRUM) was used to predict protein stability free energy change (DDG) upon single point mutation. DDG values are nearly always zero. 3 mutation classes have been described, destabilizing mutations (DDG<-0.5 Kcal/mol), stabilizing mutations (DDG>0.5 Kcal/mol) and neutral mutations (−0.5<=DDG<=0.5 Kcal/mol). The SNP A139T gave a DDG score of -1.89 suggesting it may be destabilizing to the protein.

## Discussion

Using FFPE archival tissue, 31 patients’ samples were chosen for sequencing. Several mutations were identified. *KRAS* mutations were found in 24 patients. The most prevalent was the cDNA and protein change p.G12D mutation followed by the p.G12V substitution. Both of these are known to have oncogenic effects on PDAC tumours and are associated with poor prognosis. It is expected that nearly all patients with PDACs harbour a *KRAS* mutation. This study demonstrated that only 77 percent of tumours harboured a *KRAS* mutation, this may be related to the median depth of coverage and sensitivity of the assay as opposed to these tumours being truly *KRAS* wild type. Furthermore, over 50 percent of tumours in this study harbour a *TP53* mutation. *TP53* is arguably the most important tumour suppressor gene and is present in up to 75% percent of patients with pancreatic cancer.Common variants were identified at 17: 7579472 in keeping with published data.

*PAK4* mutations in PDAC cells are rare but appear to be significant when present (Kimmelman e al., 2008). In our study, a *PAK4* frame shift mutation was found at 19:39664463 in one patient. The most interesting PAK4 finding was the presence of a A139T mutation which is also reported in the COSMIC database. Alanine-to-threonine (A to T) substitutions may deliver changes in protein confirmation (Chou and Fasman 1974), where alanine to form helices but threonine prefer to support beta-sheet structures. This is unlikely to affect the kinase domain but there are several PXXP areas on PAK4 required for protein interaction and this mutation may be present in the region of these domains. The DDG value also suggests this mutation may lead to destabilisation of the protein. Destabilisation might lead to a loss of autoinhibition and increased activity (Ha et al., 2012). We did not find any other PAKs mutated with HIGH impact in our study. In the future it would be interesting to identify copy number variation (CNV) data in this cohort. Whilst recurrent PAK mutations have not been identified, it may be that there are CNV changes in these samples.

This work demonstrates the feasibility of using FFPE samples for NGS analysis. The use of NGS for whole genome or targeted sequencing is now at the fore-front of cancer research and has filtered through to everyday clinical practice. Tumours such as melanoma, colorectal and lung cancer can now be stratified to different therapeutic pathways dependent on molecular characteristics. Whilst fresh-frozen samples are now the optimal method for storing samples for genetic analysis, there are several advantages of using FFPE samples. Firstly, it allows the molecular analysis of historic samples that are routinely stored in formalin at room temperature. Furthermore, the use of laser microdissection allows the identification of cancer tissue alongside stromal tissue, pre-cancerous lesions and normal tissue from the same individual. A limitation to this study is the small patient size.

